# A multiscale view of the Phanerozoic fossil record reveals the three major biotic transitions

**DOI:** 10.1101/866186

**Authors:** Alexis Rojas, Joaquin Calatayud, Michal Kowalewski, Magnus Neuman, Martin Rosvall

## Abstract

The hypothesis of the Great Evolutionary Faunas is a foundational concept of macroevolutionary research postulating that three global mega-assemblages have dominated Phanerozoic oceans following abrupt biotic transitions. Empirical estimates of this large-scale pattern depend on several methodological decisions and are based on approaches unable to capture multiscale dynamics of the underlying Earth-Life System. Combining a multilayer network representation of fossil data with a multilevel clustering that eliminates the subjectivity inherent to distance-based approaches, we demonstrate that Phanerozoic oceans sequentially harbored four global benthic mega-assemblages. Shifts in dominance patterns among these global marine mega-assemblages are abrupt (end-Cambrian 494 Ma; end-Permian 252 Ma) or protracted (mid-Cretaceous 129 Ma), and represent the three major biotic transitions in Earth’s history. This finding suggests that the mid-Cretaceous radiation of the so-called Modern evolutionary Fauna, concurrent with gradual ecological changes associated with the Mesozoic Marine Revolution, triggered a biotic transition comparably to the transition following the largest extinction event in the Phanerozoic. Overall, our study supports the notion that both long-term ecological changes and major geological events have played crucial roles in shaping mega-assemblages that dominated Phanerozoic oceans.

---

Sepkoski’s hypothesis of the Three Great Evolutionary Faunas that sequentially dominated Phanerozoic oceans represents a foundational concept of macroevolutionary research. This hypothesis postulates that the major groups of marine animals archived in the Phanerozoic fossil record were non-randomly distributed through time and can be grouped into Cambrian, Paleozoic, and Modern evolutionary faunas (1). Sepkoski formulated this three-phase model based on a factor analysis of family-level diversity (2), which became a framework-setting assumption in studies on the evolution of marine faunas and ecosystems (3–6), changing our view of the Phanerozoic history of life. However, because Sepkoski’s study predicts unusual volatility in the Modern evolutionary fauna starting during the mid-Cretaceous, a three-phase model fails to capture the overall diversity dynamics during long portions of the Mesozoic (7). Whether such mid-Cretaceous radiation (8) represents an intra-faunal dynamic or a biotic transition from Sepkoski’s Modern evolutionary fauna towards a neglected mid-Cretaceous-Cenozoic fauna remains unexplored.

Despite recognition that Phanerozoic marine diversity is highly structured (9), empirical estimates of the macroevolutionary pattern depend on several methodological decisions, including background assumptions, statistical threshold, hierarchical level (1, 6, 7), and the choice of input data: for example, Sepkoski’s compendia or benthic taxa from the Paleobiology Database (1, 10, 11). These limitations raise two fundamental questions: How can we identify global-scale mega-assemblage shifts without relying on critical methodological decisions? And given the underlying Earth-Life System, how should we represent the paleontological input data to accurately capture complex interdependencies? These limitations result in methodologically volatile and often inconsistent estimates of large-scale macroevolutionary structures, thus obscuring the causative drivers that underlie biotic transitions between successive global mega-assemblages. As a result, whether abrupt global perturbations, such as large bolide impacts and massive volcanic eruptions (12, 13), and long-term ecological changes (14) both operate at the higher levels of the macroevolutionary hierarchy remains unclear (14, 15).

Our understanding of the macroevolutionary dynamics of Phanerozoic life is being transformed by network-based approaches (6, 16–18). Because the input network can capture the complexity inherent to the underlying system, network analysis has become an increasingly popular alternative to the typical procedures used in almost every area of paleontological research (19–22). However, as might be expected of an emergent interdisciplinary field, methodological inconsistencies and conceptual issues in the body of network paleobiology research make it difficult to compare outcomes across studies. Also, the rapid development of the broader field of network science demands a major effort from paleobiologists working across disciplinary boundaries. Moreover, current network paleobiology studies use standard network representations based on pairwise statistics and clustering limited to a single scale of analysis (6, 18, 19, 21). That is, they use only the connection strength between nodes of geographic areas and taxa in the paleontological data and apply standard network clustering of nodes into communities, which does not capture temporal interactions between components or multiscale dynamics of the underlying Earth-Life System (23).

We employed a multilayer framework that integrates the higher-order relationships over time in the underlying paleontological data (23, 24). Specifically, our input network takes into account the temporal arrangement of fossils in the geological record, combined with multilevel hierarchical clustering (25) to test for major biotic transitions in the Phanerozoic fossil record of the benthic marine faunas (11). This multilayer network approach is transforming research on higher-order structures in both natural and social systems (26), and can help us to understand the structure and dynamics of the macroevolutionary hierarchy.

We demonstrate that Phanerozoic oceans sequentially harbored four global mega-assemblages that scale up from lower-scale biogeographic structures and shift dominance patterns across the major biotic transitions in Earth’s history. We found that abrupt global perturbations and long–term changes both played crucial roles in mega-assemblages transitions. Our study sheds light on the emergence of large-scale macroevolutionary structures (12, 27). For example, we show that biogeographic structures underlie the marine evolutionary faunas and that long-term changes controlled the shift to the modern mega-assemblage, which first emerged during the early Mesozoic but did not become dominant until the mid-Cretaceous. We also provide an integrative framework of the metazoan macroevolution for future research.

## A multilayer representation of the Earth-Life System

Standard first-order network representations, including bipartite networks and one-mode projections, overlook the temporal constraints in the underlying paleontological data and thus cannot properly capture temporal interdependencies in the Earth-Life System. A bipartite network representation of the Earth-Life System uses physical nodes for taxa and geographic areas (i.e., localities, grid cells, political units), representing its components, and weighted links between them to describe their interactions (21)(Fig. 1A). To understand the dynamics and structure of such a system, we can model the connectivity between geographic areas and taxa analyzing the trajectory of a random walker on the bipartite network, also known as network flows. T his s tandard n etwork m odel b ased on pairwise relationships in the raw data captures first-order dependencies (24): A given step on the network, for example, from a geographic area toward a taxon, depends only on the currently visited geographic area. In this representation, a random walker currently at the Coniacian Stage (∼88 Ma), for example, may move to geographic areas in the Priabonian stage (∼35 Ma) or the Santonian stage (∼85 Ma) with similar probabilities, irrespective of the previous visited stage, ignoring temporal constraints that influence the dynamics in the underlying Earth-Life System. Projecting bipartite net works into unipartite networks (28) of taxa or geographic area swashes out even more information in the paleontological data (Fig. 1B).

**Fig. 1.**
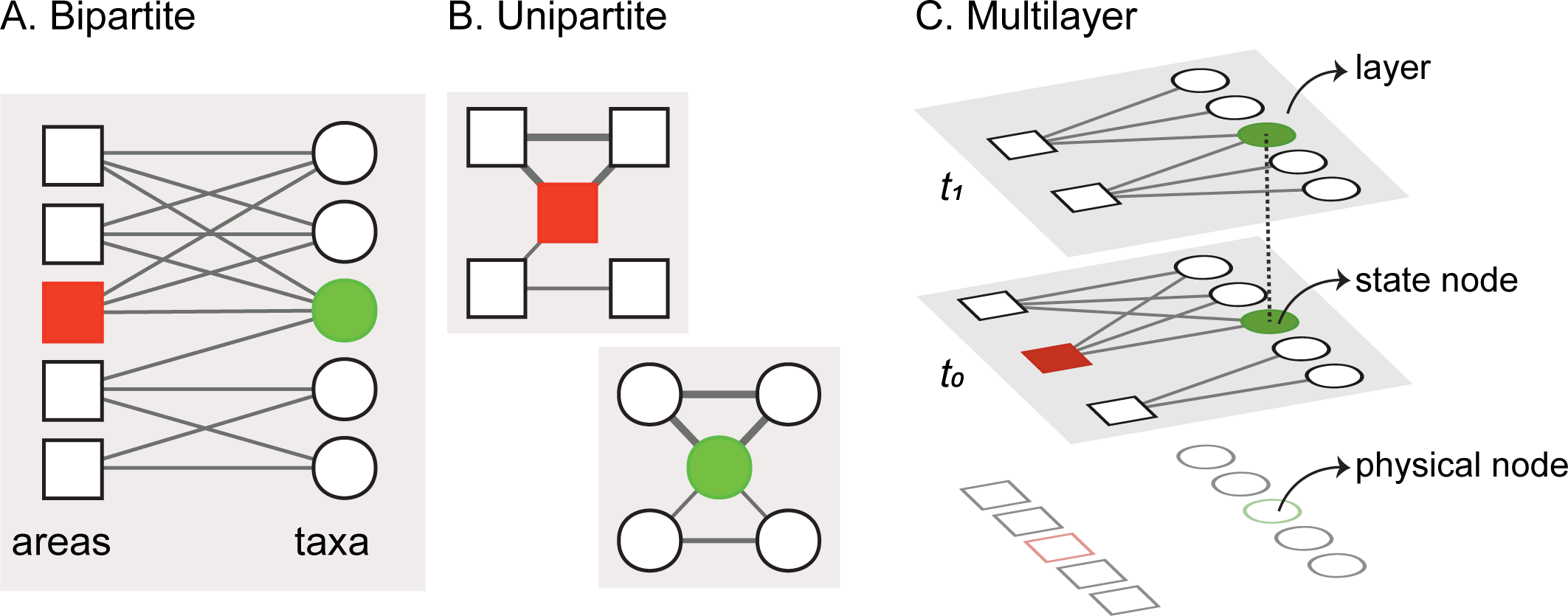
Network models used in macroevolution. A-B. Standard first-order network representations. A. Bipartite occurrence network. This representation comprises two sets of nodes that represent geographic areas and taxa (21). B. Unipartite cooccurrence networks (6, 18). These representations are weighted projections of the bipartite network onto each set of nodes. C. Higher-order multilayer representation. In this network, nodes are organized into layers representing a series of time intervals (t_0_, t_1_). The physical nodes resenting taxa are split into state nodes, with one state node per layer in which a given taxon occurs (24).

In our network representation, we account for this higher-order dynamic to reveal the multiscale organization of the Earth-Life System, which can be translated into a macroevolutionary hierarchy (27). We analyze multilayer relationships (23, 24) in the paleontological data (11) using a multilayer network framework (25, 29). Network layers represent ordered geological stages (30), and physical nodes depicting the taxa are split into state nodes (24), with one state node per each geological stage in which a given taxon occurs (Fig. 1C; Data S1). This higher-order network representation captures both the geographical and temporal aspects of the underlying Earth-Life System simultaneously.

We use the map equation multilayer framework (31), which operates directly on the assembled multilayer network and thereby preserves the higher-order interdependencies when identifying dynamical modular patterns in the data. The map equation framework consists of an objective function that measures the quality of a given network partition (32), and an efficient search algorithm that optimizes this function over different solutions (24). This algorithm provides the optimal multilevel solution for the input network, eliminating the subjectivity of distance-based approaches (6, 20). Although our input network better represents the underlying Earth-Life System compared with standard network approaches (6, 18, 21), and our clustering approach allows to capture its hierarchical modular structure, it can still be affected by numerous biases, including spatial and temporal variations in sampling effort, inequality in the rocks available for sampling, and taxonomic inconsistencies (33). We employed a parametric bootstrap to asses the potential effects of these biases on the modular structures delineated in the assembled network.

## The three major Phanerozoic biotic transitions

We found that the assembled multilayer network is best described by four significant modules at the first hierarchical level, which correspond to Phanerozoic marine mega-assemblages of highly interconnected marine benthic taxa and geographic cells (reference solution, Data S2). These large-scale modular structures characterize the underlying Earth-Life System (Fig. 2A): The Phanerozoic oceans sequentially harbored four overlapping mega-assemblages that shift dominance patterns over the three major global biotic transitions in Earth’s history, taking place at end-Cambrian (∼494 Ma), end-Permian (∼452 Ma), and mid-Cretaceous (∼129 Ma) times. The four-tier structuring of the Phanerozoic marine faunas differs from standard geological eras (Adjusted Mutual Information, AMI = 0.71), indicating that not all major biotic transitions occur at their boundaries. Although different from the three units discriminated in Sepkoski’s factor analysis (1), the classes of marine invertebrates that contribute the most to our Cambrian, Paleozoic, and combined Triassic-Cenozoic mega-assemblages match those from the hypothesis of the Three Great Evolutionary Faunas (Fig. S1). This consensus suggests that the macroevolutionary units are unlikely to represent artifacts of the factor or network analyses.

**Fig. 2.**
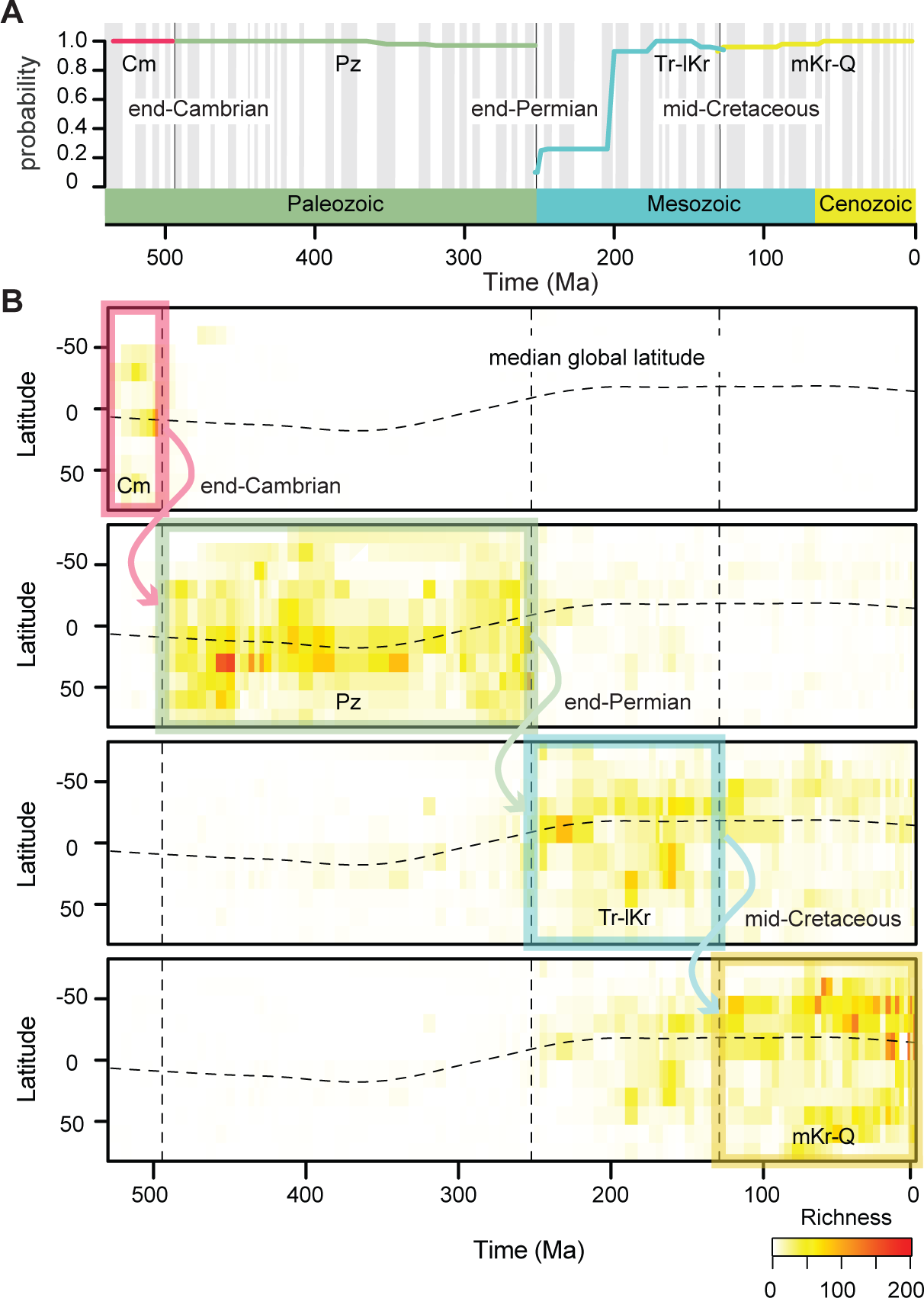
A. Large-scale modular structures in the network of Phanerozoic benthic marine faunas. The significance is the probability of retrieving a given mega-assemblage across 100 bootstrapped solutions and captures the instability of the modular structures in the assembled network after the Earth’s largest mass extinction event (34). Mega-assemblage shifts occur at the following boundaries: End-Cambrian (combined Paibian/Jiangshanian to Age10), end-Permian (Changhsingian to Induan), and mid-Cretaceous (Hauterivian to Barremian). B. Heatmaps of mega-assemblages’ genus richness across time. Heatmaps are interpolated from underlying paleontological data for 10°latitudinal bands at each geological stage. Shifts in dominance among mega-assemblages are either abrupt global perturbations (end-Cambrian and end-Paleozoic) or protracted changes with substantial spatiotemporal overlap (mid-Cretaceous). Abbreviations: Cambrian (Cm); Paleozoic (Pz); Triassic to lower Cretaceous (Tr-lKr); and mid-Cretaceous to Quaternary (mKr-Q).

The three major biotic transitions among the Phanerozoic marine mega-assemblages vary in timing (Fig. 2B) and potential causative drivers. The end-Cambrian mega-assemblage shift appears to be an abrupt transition at the base of the a uppermost Cambrian stage. However, the limited number of fossil occurrences from that interval precludes a better understanding of the transition (Materials and Methods). The end-Permian mega-assemblage shift is also abrupt. The Paleozoic and Mesozoic mega-assemblages overlap in one geological stage and share only a few taxa (Jaccard similarity index = 0.03). This biotic transition coincides with the Earth’s largest mass extinction event (34), which is considered to have caused the global shift in ocean life at that time (35). In contrast, the mid-Cretaceous mega-assemblage shift is protracted, representing a gradual shift in dominance among two mega-assemblages that share more taxa (Jaccard similarity index = 0.11) and exhibit substantial overlap in geographic space.

The protracted mid-Cretaceous mega-assemblage shift (Fig. 2B) is reminiscent of the gradual Mesozoic restructuring of the global marine ecosystems, which included changes in food-web dynamics, functional ecology of dominant taxa, and increased predation pressure (8, 36). These changes in the marine ecosystems started during the Mesozoic and continued throughout the Cenozoic (37, 38), but were particularly concentrated during the mid-Cretaceous (39). Our results suggest that such changes in the global marine ecosystems may have been responsible for the gradual emergence of modern benthic biotas. However, regardless of the specific transition mechanism, our results indicate that modern benthic biotas had already emerged during the early Mesozoic but did not become dominant until the mid-Cretaceous (∼129 Ma). In this way, the quadripartite structuring of the Phanerozoic marine fossil record captured by a multilayer network analysis couples the Three Great Evolutionary Faunas and the Mesozoic Marine Revolution hypothesis (1), which postulates the gradual diversification of the Modern evolutionary fauna during the Cretaceous (8). Sepkoski’s simulations arguably anticipated the Mesozoic transition (7) delineated in the multilayer network analysis presented here.

To evaluate the robustness of the four-phase model, we explored the landscape of alternative solutions (40). With alternative solutions obtained from parametric bootstrapping of the original network and subsequent clustering, the solution landscape shows that our four-phase model describing the Phanerozoic benthic marine faunas is highly robust to biases (Fig. 3). Alternative solutions reproduce either a four-phase model with a younger Cretaceous biotic transition, Sepkoski’s three-phase model with biotic transitions occurring at era boundaries, or a three-phase model with a mid-Cretaceous but not end-Permian biotic transition. Regardless of the number of mega-assemblages delineated, these alternative solutions demonstrate that the major biotic transitions in Earth’s history occurred across the end-Cambrian, end-Permian, and mid-Cretaceous boundaries. However, network clustering shows instability of the marine mega-assemblages at the geological stages following the Permian-Triassic boundary (Fig. 2A). The significance of the mega-assemblages drops at this boundary and then increases, likely reflecting the recovery of the benthic marine faunas and ecosystems after the Earth’s largest mass extinction event (41). Although the punctuated nature of this presumed recovery pattern should be further explored, our results indicate that the full biotic recovery from the end-Permian crisis was completed by the Early Jurassic, at which point the mega-assemblages became robust. However, our stage-level analysis does not capture the fine-scale dynamics of this biotic recovery, which is believed to have been completed during the Middle to Late Triassic (42).

**Fig. 3.**
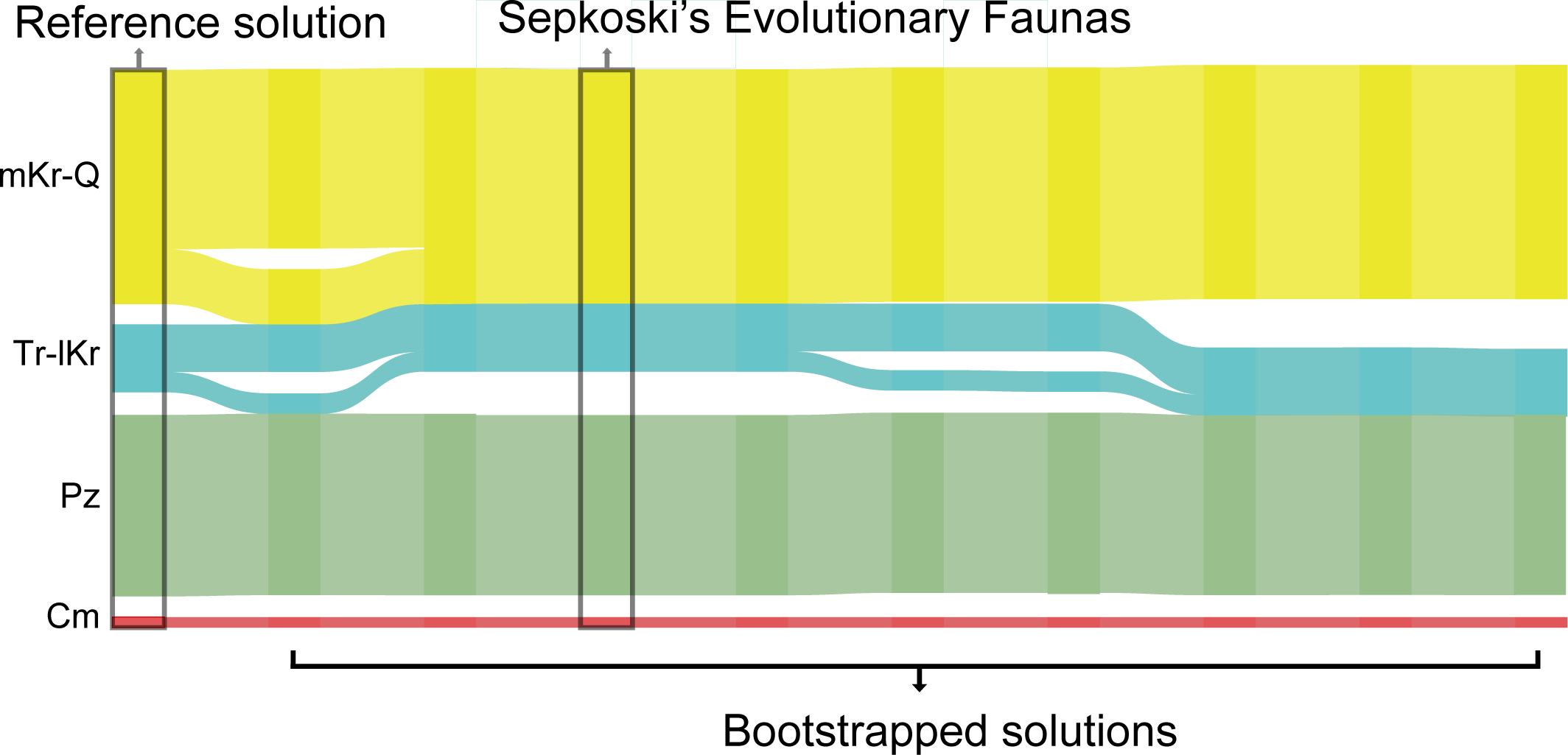
Alluvial diagram comparing our four-phase reference solution against alternative solutions obtained from bootstrapped networks. The alternative solutions represent either four-phase (Fig. 2A) or three-phase models (1). Regardless of the number of mega-assemblages delineated, these alternative solutions show that the major biotic transitions in Earth’s history occurred across the end-Cambrian, end-Permian, and mid-Cretaceous. Abbreviations: Cambrian (Cm); Paleozoic (Pz); Triassic to lower Cretaceous (Tr-lKr); and mid-Cretaceous to Quaternary (mKr-Q).

## Implications for the macroevolutionary hierarchy

We demonstrate that the Phaneozoic benthic marine faunas exhibit a hierarchically modularity in which first-level structures, representing the four Phanerozoic marine mega-assemblages, are built up from lower-level structures in a nested fashion (Fig. 4). The second-level structures underlying the four mega-assemblages represent sub-assemblages organized into time intervals that are equivalent to periods in the geological timescale (AMI = 0.83). The third- and lower-level structures underlying the mega-assemblages form geographically coherent units (21) that change over geological time. Likely due to limitations in the existing data, we were unable to map these evolutionary bioregions through the entire Phanerozoic. Nevertheless, our results demonstrate that local to regional biogeographic structures underlie the global-scale marine mega-assemblages in the macroevolutionary hierarchy. This multilevel organization of macroevolutionary units represents the large-scale spatiotemporal structure of the Phanerozoic marine diversity.

**Fig. 4.**
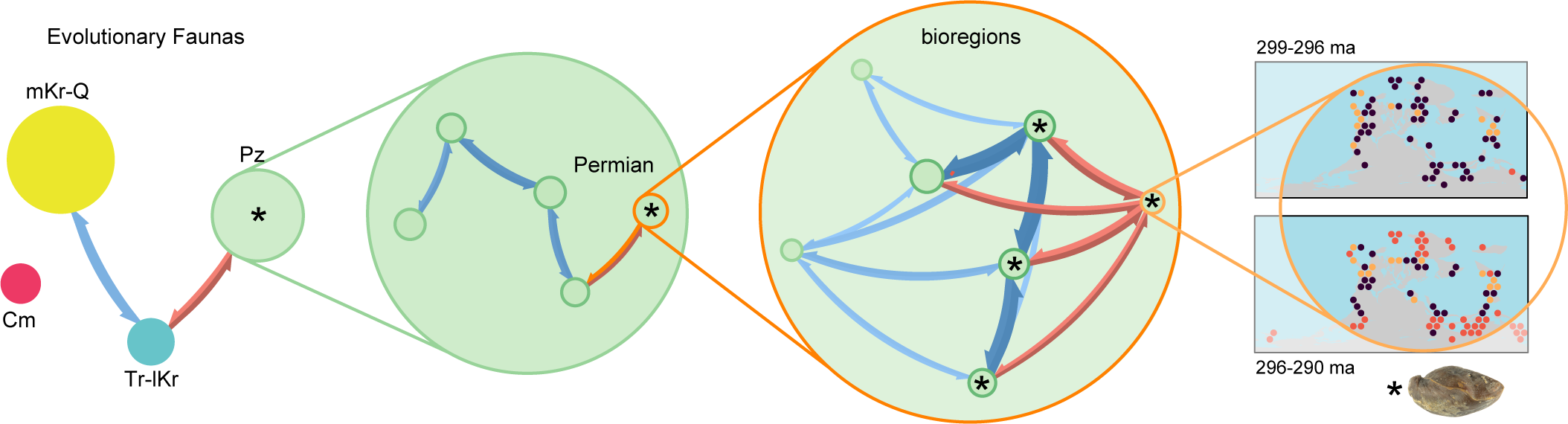
Visual representation of the nested hierarchical structures in the reference solution (Data S2). Modular structures at the higher levels of organization in this macroevolutionary hierarchy correspond to marine mega-assemblages, which are build up from lower-level entities, including sub-assemblages, evolutionary bioregions, and taxa. Elements in this representation do not represent nodes from the original network but emergent structures and their interactions.Abbreviations: Cambrian (Cm); Paleozoic (Pz); Triassic to lower Cretaceous (Tr-lKr); and mid-Cretaceous to Quaternary (mKr-Q).

**Fig. 5.**
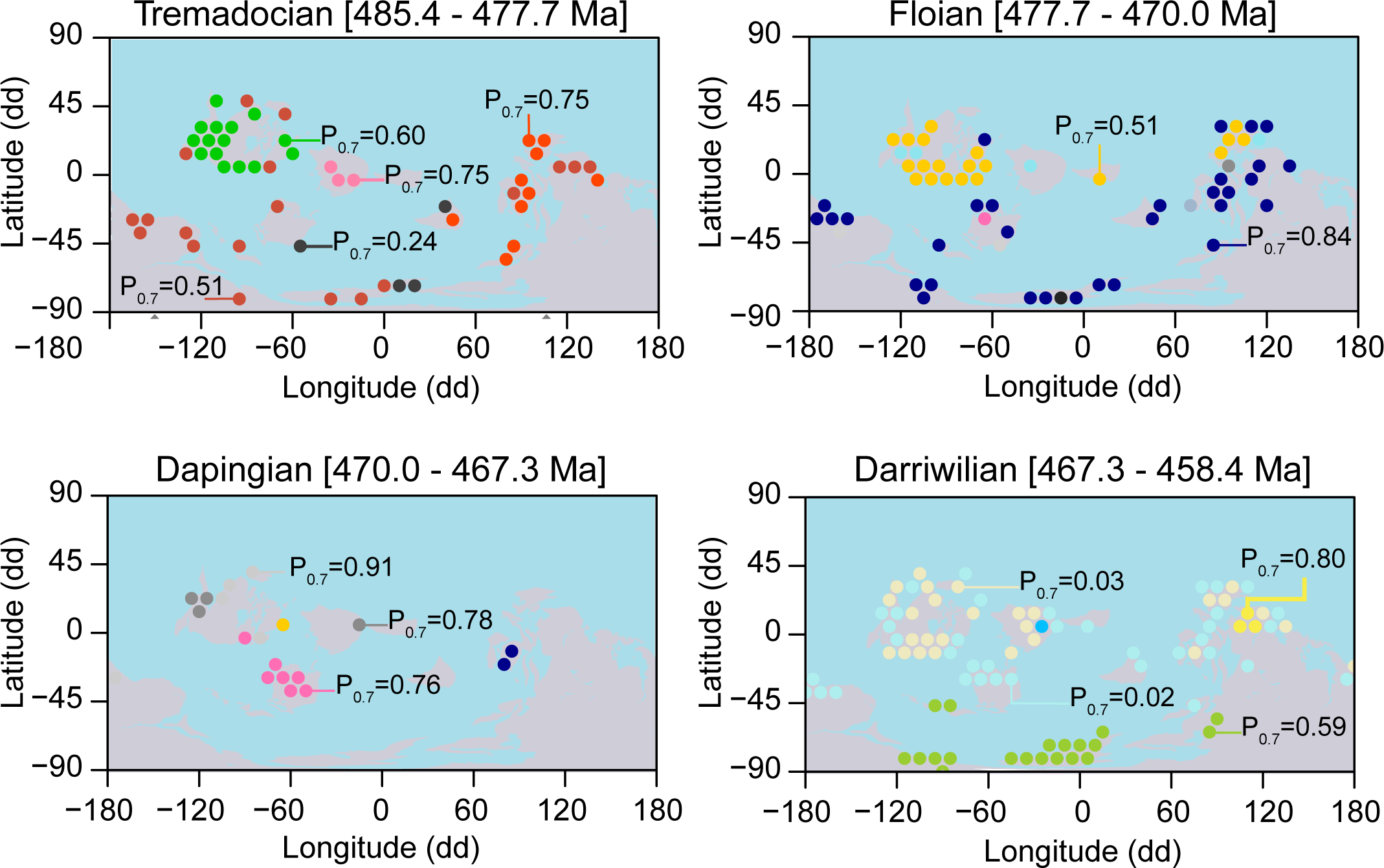
Examples of lower-level structures across geological stages. Lower-level modules form geographically coherent units underlying the Phanerozoic marine mega-assemblages. Circles represent the center of the geographic cells colored by their module affiliation (Data S2).

Without the inherent subjectivity of other approaches, our assessment of the Phanerozoic marine diversity, conducted simultaneously at different scales, can help us to comprehend the drivers and impacts most relevant at different macroevolutionary levels (12). For instance, we show that both long-term ecological interactions and global geological perturbations seem to have played a critical role in shaping the large-scale structure of the marine animals. However, some of the widely accepted major geological perturbations, including widely known global extinction events such as the Cretaceous-Tertiary extinction, control second-level but not first-level structures in this macroevolutionary hierarchy (Fig. 4). Each level of organization in the emergent macroevolutionary hierarchy is structured as a network itself and can be studied independently. Our integrative approach simultaneously quantifies both spatial and temporal aspects of the metazoan macroevolution and connects natural phenomena observed at global scales with those observed at local scales.

## Materials and Methods

### Data

We used resolved genus-level occurrences derived from the Paleobiology Database (PaleoDB) (11), which at the time of access consisted of 79,976 fossil collections with 448,335 occurrences from 18,297 genera representing the well-preserved benthic marine invertebrates (18). The PaleoDB assigns collections to paleogeographic coordinates based on their present-day geographic coordinates and age using GPlates (43). We aggregated data using paleogeographic coordinates into a regular grid of hexagons covering the Earth’s surface at each geological stage (4,906 grid cells with count > 0; inner diameter = 10° latitude-longitude) using the Hexbin R-package (http://github.com/edzer/hexbin). This binning procedure provides the symmetry of neighbors that is lacking in rectangular grids and captures the shape of geographic regions more naturally (44). The selection of an optimum grid size is a compromise between the lack of spatial resolution provided by hexagons with inner diameter => 10° and the increased number of hexagons without occurrences when shortening the inner diameter. However, recent studies have demonstrated that network analyses are robust to the shape (irregular, square and hexagonal), size (5° to 10° latitude-longitude), and coordinate system of the grid used to aggregate data (16, 45).

### Network analysis

We used the aggregated data to generate a weighted bipartite multilayer network (24), where layers represent ordered geological stages (30), and nodes represent taxa and geographic cells (21) (Fig. 1). We capture the collection-based structure of the underlying paleontological data (11) by joining taxa to geographic cells through weighted links (*w*). Specifically, for weight (*wki*) between geographic cell *k* and taxa *i*, we divided the number of collections at grid cell *k* that register taxa *i* by the total number of collections recorded at geographic cell *k*. A similar link standardization has been employed in previous studies (18, 21). We combined the last two Cambrian stages, that is, Jiangshanian Stage (494 to 489.5 Ma) and Stage 10 (489.5 to 485.4 Ma), into a single layer to account for the lack of data from the younger Stage 10 and to maintain an ordered sequence in the multilayer network framework. However, our results show that the Cambrian to Paleozoic mega assemblage shift occurred before the gap, and they are not directly related. The assembled network comprises 23,203 nodes (*n*), including 4,906 spatiotemporal grid cells and 18,297 genera, joined by 144,754 links (*m*), distributed into 99 layers (*t*) (Data S1).

We used the flow-based map equation multilayer framework with the search algorithm Infomap to cluster the assembled multilayer network (31). This high-performance clustering approach (46) allowed us to model interlayer coupling based on the intralayer information of the multilayer network using a random walker (24). The intralayer link structure represents the geographic constraints on network flows at a given geological stage in Earth’s history, and the interlayer link structure represents the temporal ordering of those stages. In this neighborhood flow coupling, a random walker within a given layer moves between taxa and geographic cells guided by the weighted intralayer links with probability (1-*r*), and it moves guided by the weighted links in both the current and adjacent layers with a probability *r*. Consequently, the random walker tends to spend extended times in multilayer modules of strongly connected taxa and geographic cells that correspond to Phanerozoic marine mega-assemblages. Following the methodology of previous studies, we used the relax rate *r* =0.25 which is large enough to enable interlayer temporal dependencies but small enough to preserve intralayer geographic information (47). We tested the robustness to the selected relax rate by clustering the assembled network for a range of relax rates and compared each solution to the solution for *r* =0.25 using Jaccard Similarity. We obtained the reference solution (Data S2) using the assembled network and the following Infomap arguments: `-N 200 -i multilayer –multilayer-relax-rate` `0.25 –multilayer-relax-limit 1`. The relax limit is the number of adjacent layers in each direction to which a random walker can move; a value of 1 enables temporal ordering of geological stages in the multilayer framework.

### Robustness analysis

We employed a parametric bootstrap to estimate the significance of the four Phanerozoic megaassemblages in the reference solution. This standard approach accounts for the uncertainty in the weighted links connecting taxa to geographic cells due to numerous biases (33). We resampled taxon occurrence using a truncated Poisson distribution with mean equal to the number of taxon occurrences. The truncated distribution has all probability mass between one and the total number of collections in the grid cell, thus avoiding false negatives. We obtained the resampled link weight by dividing the sampled number by the total number of recorded collections. Using Infomap with the arguments detailed above, we clustered these bootstrapped networks and compared the results against the reference solution. Specifically, for each reference module, we computed the proportion of bootstrapped partitions where we could find a module with Jaccard similarity higher than 0.5 (P_05_) and 0.7 (P_07_) (Tables S1-S2). We also computed the average probability (median) of belonging to a supermodule for nodes of the same layer (Fig. 2A). This procedure for estimating module significance is detailed in ref. (40).

## ACKNOWLEDGMENTS

We thank the contributors to the Paleobiology Database who collected data. We thank S. Finnegan and D. Edler for useful discussions, and R. Nawrot for helpful comments on an early version of the manuscript. A.R. was supported by the Olle Engkvist Byggmästare Foundation, J.C. by the Carl Trygger Foundation, and M.R. by the Swedish Research Council, grant 2016-00796.

---

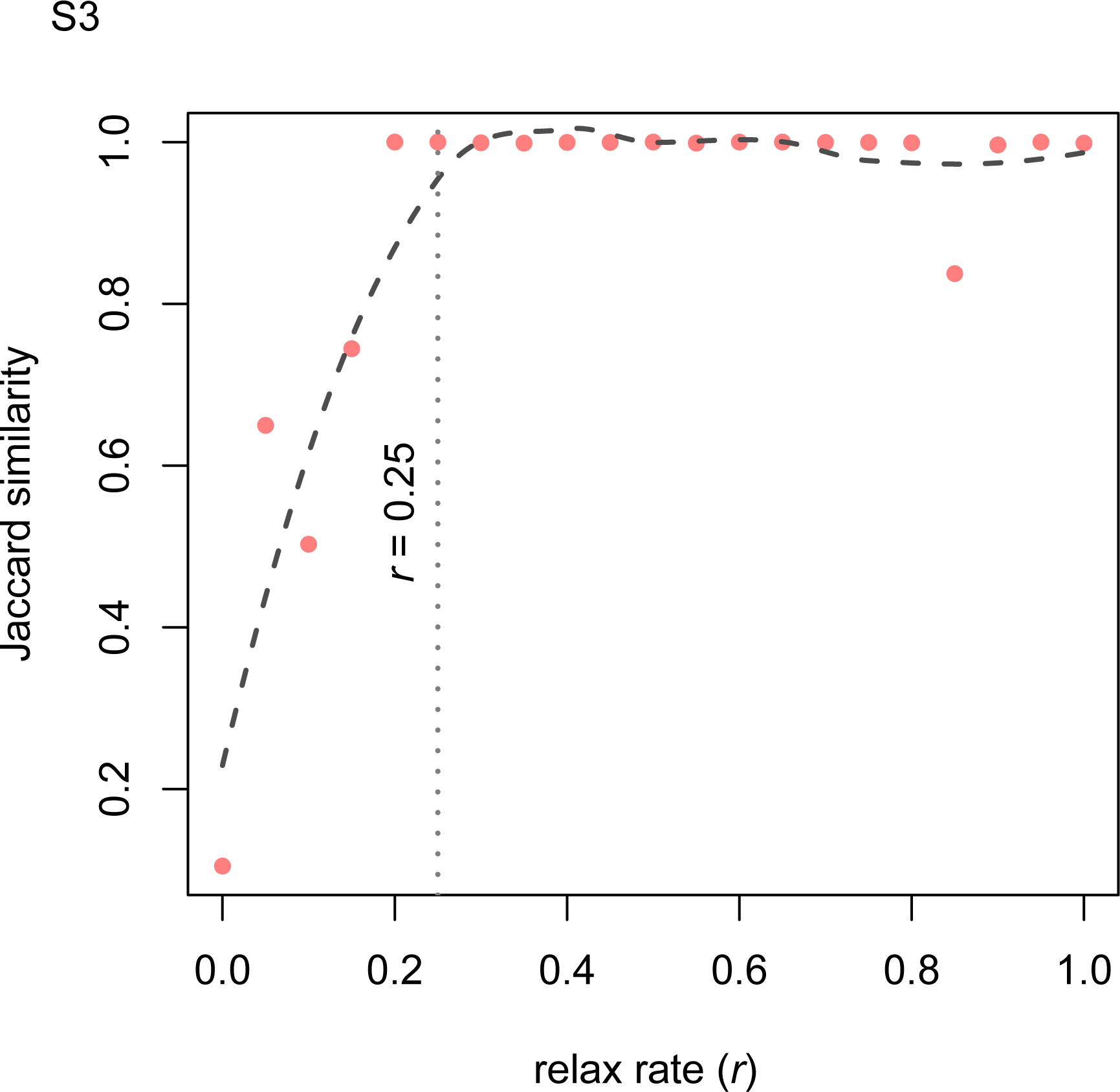

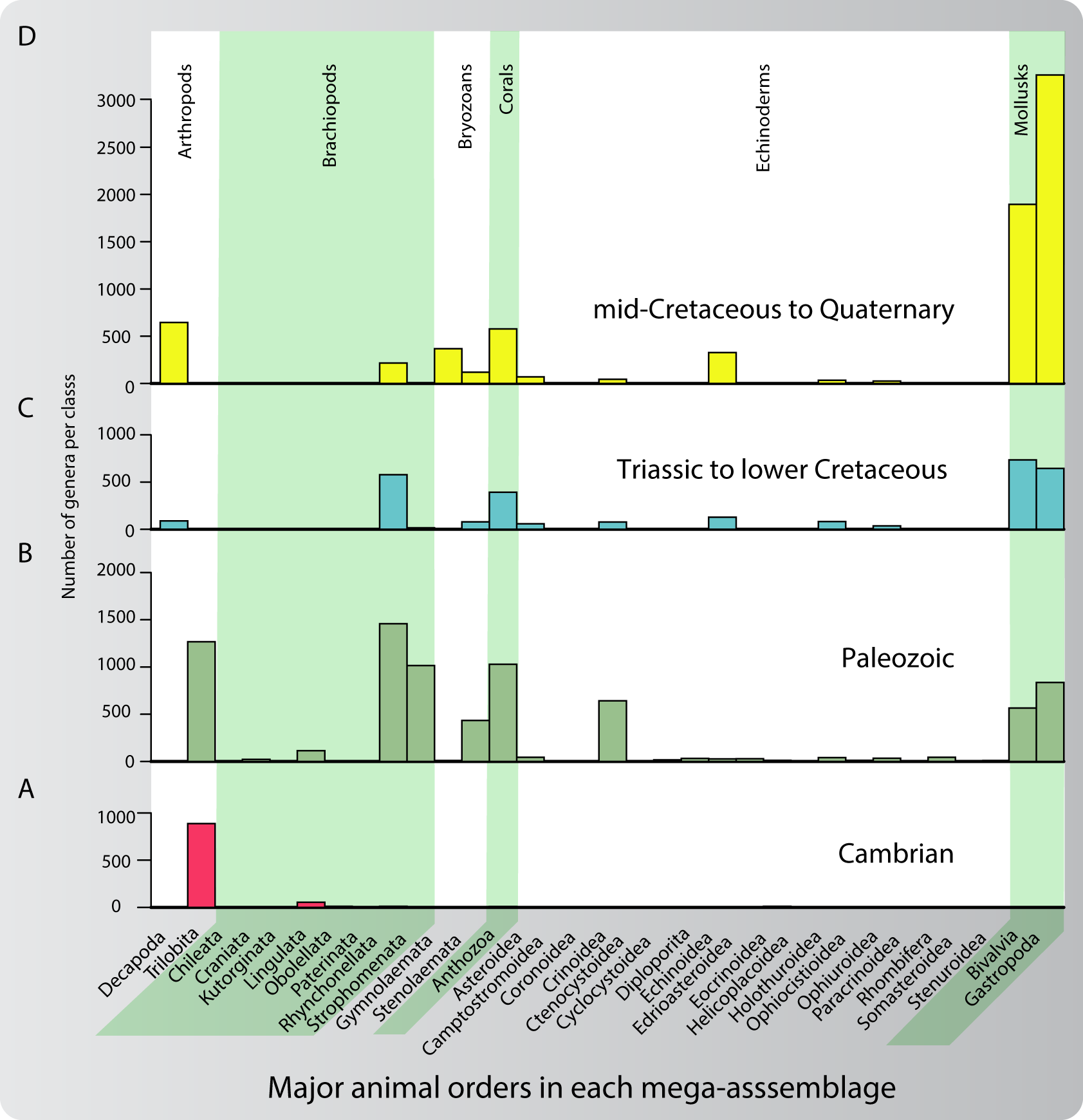

## References

1. Sepkoski JJ (1981) A factor analytic description of the Phanerozoic marine fossil record. Paleobiology 7(01):36–53.

2. Sepkoski JJ (1984) A kinetic model of Phanerozoic taxonomic diversity. III. Post-Paleozoic families and mass extinctions. Paleobiology 10(2):246–267.

3. Peters SE (2004) Relative abundance of Sepkoski’s evolutionary faunas in Cambrian-Ordovician deep subtidal environments in North America. Paleobiology 30(4):543–560.

4. Meroi Arcerito FR, Halpern K, Balseiro D, Waisfeld B (2017) Tempo and mode in the replacement of trilobite evolutionary faunas from the Cordillera Oriental basin (Northwestern Argentina). Comptes Rendus Palevol 16(8):821–831.

5. Brayard A, et al. (2017) Unexpected Early Triassic marine ecosystem and the rise of the Modern evolutionary fauna. Science Advances 3(2):e1602159.

6. Muscente AD, et al. (2018) Quantifying ecological impacts of mass extinctions with network analysis of fossil communities. Proceedings of the National Academy of Sciences 115(20):5217–5222.

7. Alroy J (2004) Are Sepkoski’s evolutionary faunas dynamically coherent? Evolutionary Ecology Research 6(1):1–32.

8. Vermeij GJ (1977) The Mesozoic marine revolution: evidence from snails, predators and grazers. Paleobiology 3(3):245–258.

9. Hofmann R, Tietje M, Aberhan M (2019) Diversity partitioning in Phanerozoic benthic marine communities. Proceedings of the National Academy of Sciences 116(1):79–83.

10. Sepkoski JJ (1996) Patterns of Phanerozoic Extinction: a Perspective from Global Data Bases in Global Events and Event Stratigraphy in the Phanerozoic, ed. Walliser OH. (Springer Berlin Heidelberg, Berlin, Heidelberg), pp. 35–51.

11. Peters SE, McClennen M (2016) The Paleobiology Database application programming interface. Paleobiology 42(01):1–7.

12. Myers CE, Saupe EE (2013) A macroevolutionary expansion of the modern synthesis and the importance of extrinsic abiotic factors. Palaeontology 56(6):1179–1198.

13. Hull PM, et al. (2020) On impact and volcanism across the Cretaceous-Paleogene boundary. Science 367(6475):266–272.

14. Voje KL, Holen ØH, Liow LH, Stenseth NC (2015) The role of biotic forces in driving macroevolution: beyond the Red Queen. Proceedings of the Royal Society B: Biological Sciences 282(1808):20150186.

15. Benton MJ (2009) The Red Queen and the Court Jester: Species Diversity and the Role of Biotic and Abiotic Factors Through Time. Science 323(5915):728–732.

16. Vilhena DA, et al. (2013) Bivalve network reveals latitudinal selectivity gradient at the end-Cretaceous mass extinction. Scientific Reports 3.

17. Dunhill AM, Bestwick J, Narey H, Sciberras J (2016) Dinosaur biogeographical structure and Mesozoic continental fragmentation: a network-based approach. Journal of Biogeography.

18. Kocsis AT, Reddin CJ, Kiessling W (2018) The biogeographical imprint of mass extinctions. Proceedings of the Royal Society B: Biological Sciences 285(1878):20180232.

19. Vilhena DA, Antonelli A (2015) A network approach for identifying and delimiting biogeographical regions. Nature Communications 6:6848.

20. Kiel S (2016) A biogeographic network reveals evolutionary links between deep-sea hydrothermal vent and methane seep faunas. Proceedings of the Royal Society B: Biological Sciences 283(1844):20162337.

21. Rojas A, Patarroyo P, Mao L, Bengtson P, Kowalewski M (2017) Global biogeography of Albian ammonoids: A network-based approach. Geology 45(7):659–662.

22. Muscente AD, et al. (2019) Ediacaran biozones identified with network analysis provide evidence for pulsed extinctions of early complex life. Nature Communications 10(1):911.

23. Xu J, Wickramarathne TL, Chawla NV (2016) Representing higher-order dependencies in networks. Science Advances 2(5):e1600028.

24. Edler D, Bohlin L, Rosvall a (2017) Mapping Higher-Order Network Flows in Memory and Multilayer Networks with Infomap. Algorithms 10(4):112.

25. De Domenico M, Lancichinetti A, Arenas A, Rosvall M (2015) Identifying modular flows on multilayer networks reveals highly overlapping organization in interconnected systems. Physical Review X 5(1):011027.

26. Siyari P, Dilkina B, Dovrolis C (2019) Emergence and Evolution of Hierarchical Structure in Complex Systems in Dynamics On and Of Complex Networks III, eds. Ghanbarnejad F, Saha Roy R, Karimi F, Delvenne JC, Mitra B. (Springer International Publishing, Cham), pp. 23–62.

27. Jablonski D (2017) Approaches to Macroevolution: 1. General Concepts and Origin of Variation. Evolutionary Biology 44(4):427–450.

28. Zhou T, Ren J, Medo M, Zhang YC (2007) Bipartite network projection and personal recommendation. Physical Review E 76(4):046115.

29. Mucha PJ, Richardson T, Macon K, Porter MA, Onnela JP (2010) Community structure in time-dependent, multiscale, and multiplex networks. Science 328(5980):876–878.

30. Gradstein FM, Ogg JG, Smith AG (2004) A geologic time scale 2004. (Cambridge University Press, Cambridge, UK; New York).

31. Edler D, Eriksson A, Rosvall M (2019) The Infomap Software Package.

32. Rosvall M, Bergstrom CT (2008) Maps of random walks on complex networks reveal community structure. Proceedings of the National Academy of Sciences 105(4):1118–1123.

33. Smith AB (2007) Marine diversity through the Phanerozoic: problems and prospects. Journal of the Geological Society 164(4):731–745.

34. Song H, Wignall PB, Dunhill AM (2018) Decoupled taxonomic and ecological recoveries from the Permo-Triassic extinction. Science Advances 4(10):eaat5091.

35. Penn JL, Deutsch C, Payne JL, Sperling EA (2018) Temperature-dependent hypoxia explains biogeography and severity of end-Permian marine mass extinction. Science 362(6419):eaat1327.

36. Fraaije RH, van Bakel BW, W.M. Jagt J, Andrade Viegas P (2018) The rise of a novel, plankton-based marine ecosystem during the Mesozoic: a bottom-up model to explain new higher-tier invertebrate morphotypes. Boletín de la Sociedad Geológica Mexicana 70(1):187–200.

37. Leckie RM, Bralower TJ, Cashman R (2002) Oceanic anoxic events and plankton evolution: Biotic response to tectonic forcing during the mid-Cretaceous. Paleoceanography 17(3):13–1–13–29.

38. Knoll AH, Follows MJ (2016) A bottom-up perspective on ecosystem change in Mesozoic oceans. Proceedings of the Royal Society B: Biological Sciences 283(1841):20161755.

39. Knoll AH (2003) Biomineralization and Evolutionary History. Reviews in Mineralogy and Geochemistry 54(1):329–356.

40. Calatayud J, Bernardo-Madrid R, Neuman M, Rojas A, Rosvall M (2019) Exploring the solution landscape enables more reliable network community detection. Physical Review E 100(5):052308.

41. Penn JL, Deutsch C, Payne JL, Sperling EA (2018) Temperature-dependent hypoxia explains biogeography and severity of end-Permian marine mass extinction. Science 362(6419):eaat1327.

42. Chen ZQ, Benton MJ (2012) The timing and pattern of biotic recovery following the end-Permian mass extinction. Nature Geoscience 5(6):375–383.

43. Müller RD, et al. (2018) GPlates: Building a Virtual Earth Through Deep Time. Geochemistry, Geophysics, Geosystems 19(7):2243–2261.

44. Birch CP, Oom SP, Beecham JA (2007) Rectangular and hexagonal grids used for observation, experiment and simulation in ecology. Ecological Modelling 206(3-4):347–359.

45. Costello MJ, et al. (2017) Marine biogeographic realms and species endemicity. Nature Communications 8(1):1057.

46. Lancichinetti A, Fortunato S (2009) Community detection algorithms: A comparative analysis. Physical Review E 80(5):056117.

47. Aslak U, Rosvall M, Lehmann S (2018) Constrained information flows in temporal networks reveal intermittent communities. Physical Review E 97(6):062312.

